# Simulation of neurotransmitter release and its imaging by fluorescent sensors

**DOI:** 10.64898/2026.03.23.707923

**Authors:** Juliana Gretz, Jennifer M. Mohr, Bjoern F. Hill, Valeriia Andreeva, Luise Erpenbeck, Sebastian Kruss

## Abstract

Cells release signaling molecules such as neurotransmitters that diffuse through the extracellular space and bind to receptors. These signaling molecules can be detected by fluorescent sensors/probes to provide images of the signaling process. Such images are not equivalent to a concentration because diffusion and sensor kinetics affect (convolute) them. Therefore, computational approaches are necessary to disentangle these contributions and allow interpretation of fluorescent sensor-based images. Here, we present a kinetic Monte Carlo framework (FLuorescence Imaging Kinetic Simulation, FLIKS) that simulates signaling molecules undergoing cellular release, stochastic diffusion and reversible binding to sensors in realistic cellular (2D or 3D) geometries. We apply it to model neurotransmitter (dopamine) release in synaptic clefts and for paracrine signaling by immune cells. We also show how sensor location, sensor kinetics and release location affect fluorescence images. For example, we show how sensor sensitivity depends on the distance from the synaptic cleft and changes when dopamine transporters (DAT) clear dopamine. The approach also allows to compare the performance of membrane bound (genetically encoded) sensors versus artificial sensors such as nanosensors placed outside under or around the cells. As an example, we also demonstrate how the images of catecholamine release by immune cells can be modeled and compared to experimental data to better understand the release pattern. This framework provides a quantitative basis for analyzing and interpreting fluorescent sensor imaging data.

## Introduction

Intercellular communication in biological systems relies on the precise spatiotemporal control of molecular signaling.^1^ First, signaling molecules are released. Next, these molecules diffuse through extracellular space until they encounter a receptor or a transporter. The spatial arrangement of release sites, diffusion barriers, receptors as well as binding kinetics determines how information is processed and integrated by the cells. Insights into these parameters are key to understanding how cells coordinate their communication. These principles apply across various biological compartments such as synaptic neurotransmission^2^, paracrine signaling^3^, and immune cell-cell interactions^4^.

One of the most important neurochemicals is dopamine, which acts as a neuromodulator and therefore diffusion through the tissue contributes to its biological function.^5^ Dopamine release in neurons is linked to reward control and disorders of the dopaminergic system are associated with Parkinson’s disease, depression or Alzheimer’s disease.^6–9^ Beyond the nervous system, dopamine plays immunomodulatory roles through receptors on neutrophils, lymphocytes, monocytes, and mast cells, regulating inflammatory signaling and effector functions.^10–13^ Despite its importance, key questions such as the role of volume transmission^14^ or co-release are not fully understood^15,16^ These open questions motivate the development of novel sensing and imaging technologies with high spatial, temporal and chemical resolution.

Genetically encoded fluorescent (protein-based) biosensors such as dLight^17^ and GRAB_DA_^18^ combine a dopamine receptor binding domain and a fluorescent protein enabling real-time dopamine imaging *in vivo* with cell-type specificity^17^. Analogous sensors exist for glutamate (iGluSnFR)^19^ or norepinephrine (GRAB_NE_)^20^, with multiple variants per family covering distinct concentration ranges. For example, dLight1.1 (K_D_ ∼ 330 nM) and dLight1.3b (K_D_ ∼ 2.4 µM) can resolve distinct tonic and phasic release reglugimes.^17^ However, these sensors operate only in a certain spectral range and require efficient transfection.

An alternative are artificial sensors. Here, single-walled carbon nanotubes (SWCNTs) have shown great potential as near infrared (NIR) fluorescent probes.^21^ SWCNT-based sensors are reported for signaling molecules such as catecholamines including dopamine^22^, serotonin^23,24^, oxytocin^25^ or reactive oxygen species^26,27^. These sensors can also be designed for other biomolecule classes such as cancer markers^28^ , fibrinogen^29^ or hydrolase enzyme activity in soil^30^. They are photostable^31^ and work in complex biological media.^22,32,33^ With a layer of these sensors positioned beneath cultured cells, cellular dopamine efflux at subcellular resolution was captured, revealing heterogeneous release hotspots and extracellular concentration gradients.^32,34–36^ Ratiometric dual-emission approaches further enhance detection robustness.^34^ By single-molecule experiments the typical rate constants of such sensors were determined and it was shown that they are DNA sequence dependent.^37^ Typical sensors are best described by two first order kinetics with rate constants of 𝑘_𝑜𝑓𝑓,𝑓𝑎𝑠𝑡_ = 0.40 − 0.71 𝑠^−1^ and 𝑘_𝑜𝑓𝑓,𝑠𝑙𝑜𝑤_ = 0.01 − 0.10 𝑠^−1^ and 𝑘_𝑜𝑛_ = 1 · 10^4^ − 8 · 10^4^ 𝑀^−1^𝑠^−1^, depending on DNA sequence and analyte.^37^

Despite advances in optical, electrochemical, and nanoparticle-based sensors, direct experimental access to these spatiotemporal processes remains limited.^38^ Sensor measurements and imaging provides a representation of the analyte concentration but it is convoluted by sensor kinetics^39^, geometry, and position.^27,40^ Binding kinetics can be interpreted as filters that blur the concentration.^41–43^ Furthermore, the heterogeneous extracellular space generates concentration gradients that contribute to the biological complexity.^44^ Understanding how diffusion shapes the concentration profile measured by sensors is essential for interpreting signaling dynamics. Even though dopamine is a very important case these principles apply broadly to signaling molecules across diverse biological contexts.

Experimental limitations have an additional impact on the fluorescence readout such as the diffraction limit of light or the distribution of the sensors as well as their impact on the concentration profile. These experimental constraints motivate computational approaches that systematically predict how diffusion, geometry, sensor location and binding kinetics shape the spatiotemporal fluorescence images that represent chemical signaling.

In this work, we provide a solution for this challenge (**Figure 1a**). We simulate how analyte diffusion and binding kinetics shape the images of extracellular signaling captured by fluorescent sensors (“Fluorescent Sensor Imaging Kinetic Simulation”, FLIKS). We examine how release site geometries, diffusion barriers, transporters and sensor location influence the obtainable fluorescence images/traces in space and time. This enables us to identify conditions that allow to distinguish biological scenarios. It provides a framework to understand and interpret quantitative data from fluorescent sensor imaging, for example of neurotransmitter sensors.

**Figure 1:**
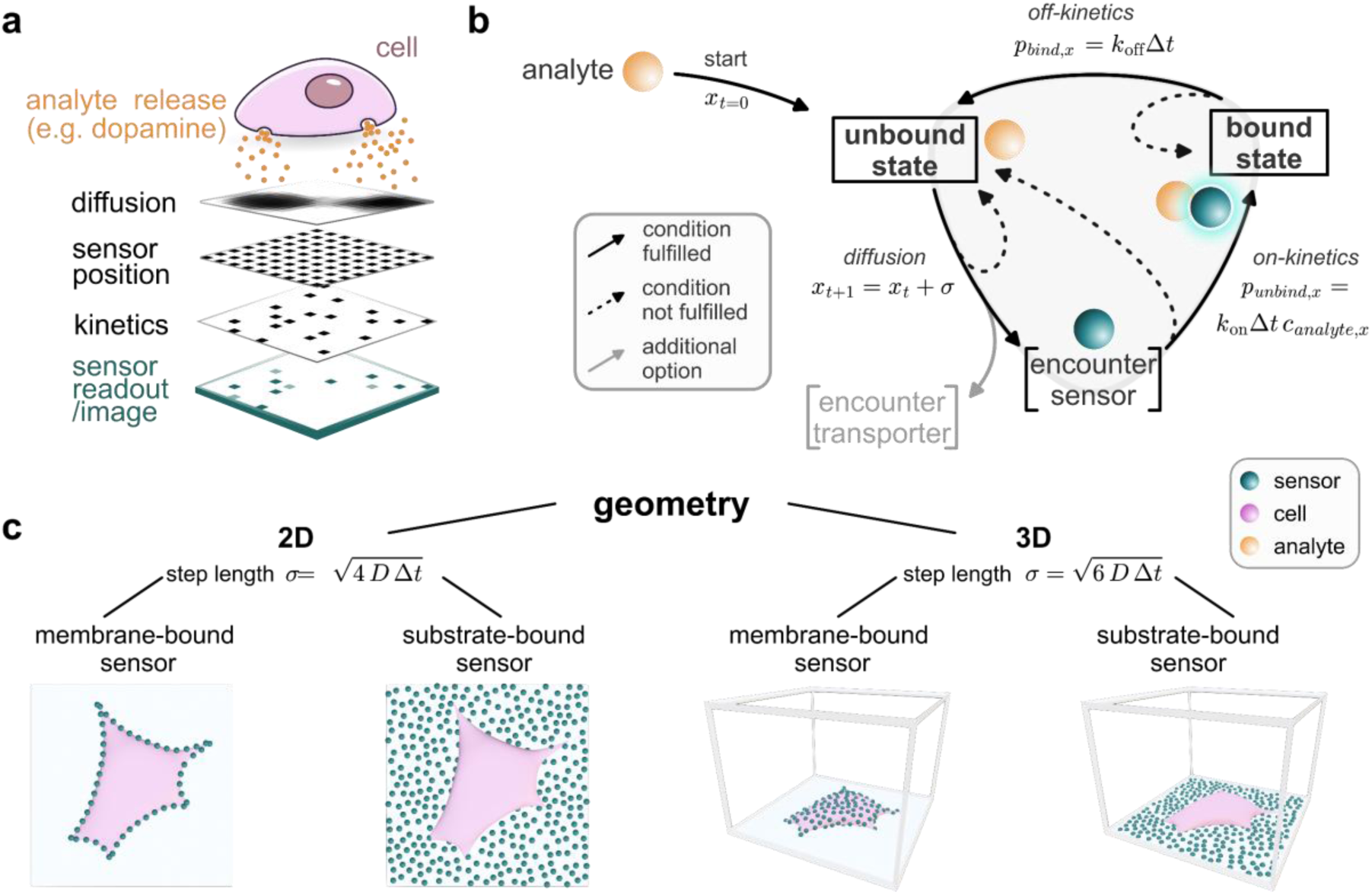
Framework to simulate how extracellular (neurotransmitter) signaling is captured by imaging of sensors (Fluorescent Sensor Imaging Kinetic Simulation, FLIKS). a) Overview of factors affecting an image (=sensor readout) e.g. from an exocytosis event. Analyte release (orange) from cells (pink) is a biological event, but sensor readout (green) depends on diffusion, sensor position and binding kinetics. b) Schematic of the stochastic simulation algorithm. Analyte position 𝑥 at a given time 𝑡 updates as 𝑥_𝑡+1_ = 𝑥_𝑡_ + 𝜎. Here, 𝜎 is the length of a step. Analytes transition between unbound and bound states with reaction kinetics described by the binding probabilities 𝑝_𝑢𝑛𝑏𝑖𝑛𝑑,𝑥_ and 𝑝_𝑏𝑖𝑛𝑑,𝑥_ with rate constants 𝑘_𝑜𝑓𝑓_ and 𝑘_𝑜𝑛_. 𝛥 𝑡 is the (smallest) simulation time step and 𝑐_𝑎𝑛𝑎𝑙𝑦𝑡𝑒,𝑥_ is the analyte concentration at position 𝑥. c) Implementation options for different geometries and positions of sensors. Membrane-bound is the expected standard case for expressed protein-based fluorescent sensors. Non-protein-based sensors such as nanosensors can be placed on cell surfaces or the substrate (e.g. the glass under a cell in cell culture).

## Results and Discussion

Our simulation is based on a kinetic Monte Carlo (kMC) simulation (**Figure 1b**). We named it Fluorescent Sensor Imaging Kinetic Simulation (FLIKS). It simulates individual analyte molecules undergoing diffusive motion with stochastic binding and unbinding events to sensors, receptors or transporters and is governed by kinetic rate constants 𝑘_𝑜𝑛_ and 𝑘_𝑜𝑓𝑓_. It can model planar sensor geometries with sensors either on substrates such as glass or on the membrane both in 2D or 3D (**Figure 1c**). For details of the simulation, we refer to the materials and methods section and the corresponding python scripts.

To understand how sensor localization/positioning affects the images one obtains from fluorescent sensors, we simulated neurotransmitter release (here dopamine) in a realistic synaptic geometry (Figure 2a).^45^ Note that the concept is generic and not limited to specific cells or signaling molecules. We focused on three scenarios that match the typical location of different sensor technologies a) expression of the sensor on the cell membrane b) expression or positioning inside the synaptic cleft c) placement outside the synaptic cleft in the extracellular space for example when artificial sensors are immobilized on a substrate.

**Figure 2:**
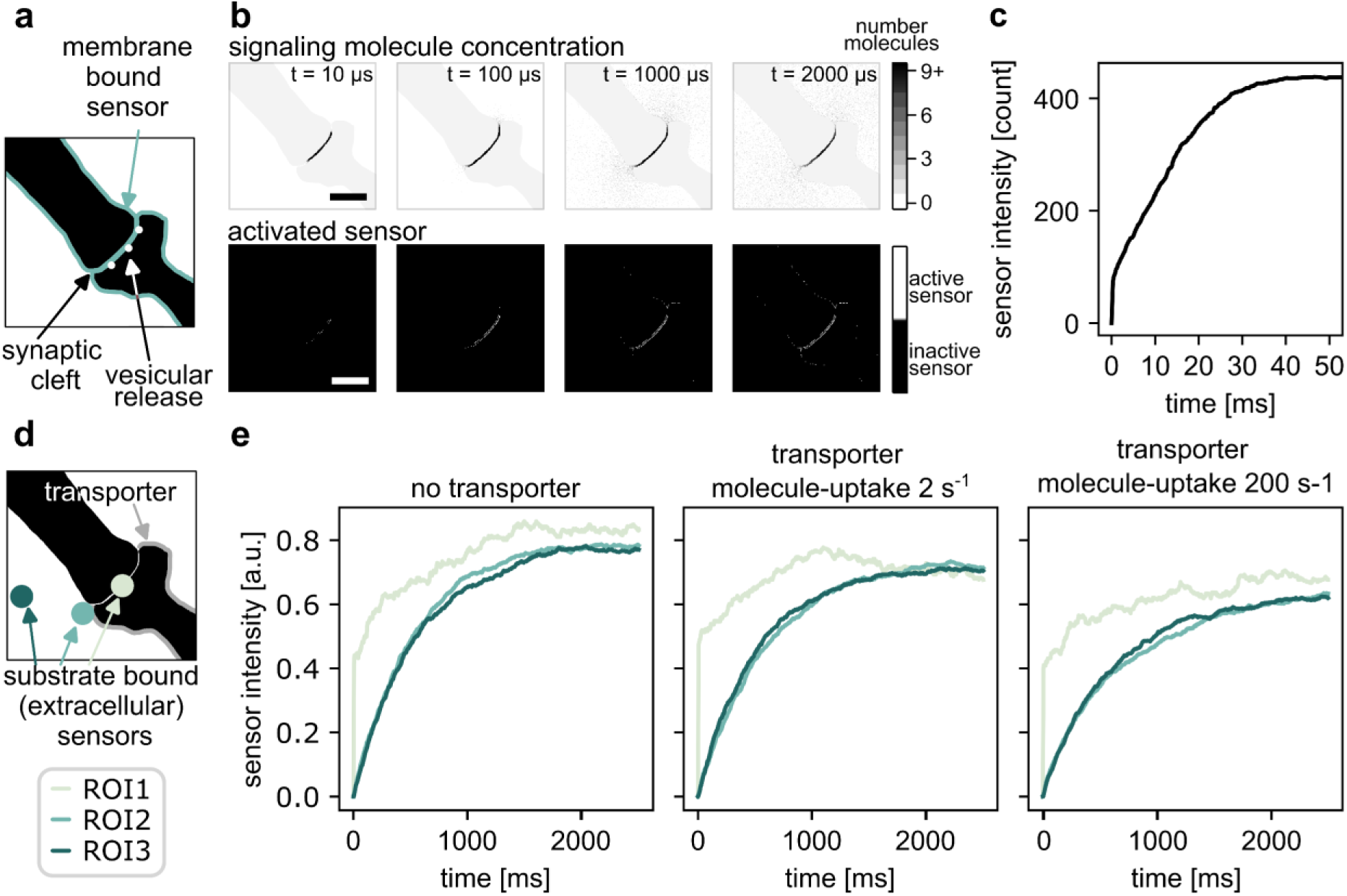
Spatial and temporal dynamics of dopamine imaging in and around a synapse. a) Simulation geometry of a typical synaptic cleft^45^ (black) with membrane-bound sensors (cyan). **b)** Spatiotemporal evolution of dopamine concentration (top row) and activated sensor (= fluorescence change by analyte binding) distribution (bottom row) at t = 10, 100, 1000 and 2000 µs after release. Scale bars = 1 µm. **c)** Time resolved course of analyte-sensor-binding from (b): total bound molecules on total sensor surface over time. Note that saturation corresponds to an equilibrium in the simulation box (see video M1, and table T1 for details). **d)** Exemplary sensor configuration for assessing the effect of uptake via a dopamine transporter (DAT). Three sensor patches (s1-s3, cyan, diameter = 0,5 µm) are positioned at varying distances from the release site. DAT (orange) is positioned on the membrane of the presynapse. The vesicular release is similar to a. **e)** Sensor intensity over 2.5 seconds with different transporter uptake rates: without transporter, with a transporter taking up 2 molecules per second (DAT^46^) and with a transporter taking up 200 molecules per second. Videos: Video S1-S4; Parameters: Table S1-S2.

As an initial scenario, sensors were positioned on the membrane either outside, within the synaptic cleft or in the extracellular space (Figure 2a). These sensors were modeled according to the expected kinetics of the genetically encoded sensor nLight.^47^ Following the fusion of exemplary (here three) vesicles, dopamine diffused through the synaptic cleft and subsequently spilled over into the extra synaptic space (Figure 2b, upper panel). The membrane-bound sensor was activated according to the local dopamine concentration and kinetics (Figure 2b, lower panel). First binding events occurred within microseconds, on the post- and presynaptic membrane (Figure 2c). After 50 ms, dopamine started to spill out of the cleft and activated sensors on the extracellular membrane. After around 30 ms, the number of bound molecules reached a plateau in this 4x4 µm simulated area. The plateau can be explained because the analyte cannot diffuse out of the simulation box. However, one can expect that in brain tissue diffusion is also to a certain degree confined.^48^

A limitation of the simulation involves the handling of diffusion boundaries. Molecules that would cross a boundary are currently retained at their current position rather than reflected. This can artificially slow down diffusion. A more accurate approach was explored but discarded due to excessive computational cost. To compensate for this bias, time increments were kept short (10 µs).

To investigate how sensor responses depend on their positioning relative to the release site, three exemplary sensors patches (s1-s3, diameter 500 nm) were positioned at increasing distance from the release site (Figure 2d). It reflects the expected size of a resolution limited spot for a NIR fluorescent nanosensor with emission wavelength of around 1000 nm. For these sensors kinetic parameters of functionalized single-walled carbon nanotube (SWCNT) sensors on a glass surface were used.^37^ Unlike the idealized scenario in panel b, sensors cannot be perfectly arranged in a pixel-like grid. Additionally, the spatial resolution is limited by the diffraction limit of fluorescence microscopy.

Without dopamine transporter (DAT) activity, all three sensors eventually reached steady-state binding levels after approximately 2 seconds (Figure 2e, left panel). However, the temporal dynamics differed between sensors. Sensor s1 was positioned closest to the release site. It showed a rapid initial burst of binding, comparable to the membrane-bound sensor (Figure 2c). Sensors s2 and s3 were located further away. Their response was delayed and dampened.

In physiological conditions, DAT actively clears dopamine from the extracellular space. DATs are membrane proteins on presynaptic neurons that transport dopamine back into the cell. ^49^ Critically, DATs are localized primarily outside the synapse rather than within the synaptic cleft.^50^ Therefore, we also checked the impact of DAT activity (Figure 2e, right panel). DAT activity reduced activation of both sensors positioned outside of the synaptic cleft (s2 and s3). Sensor s2 was most affected, as it was positioned close to the transporter. However, the transporter activity did not have a big impact in sensor activation and concentration, which shows that dopamine mainly acts through volume transmission. Another scenario with a 100x higher uptake rate shows a stronger decrease in intensity. These findings illustrate how transporter uptake rate, volume transmission and the structure of the extracellular space could affect signaling.

To examine how the position of analyte release affects sensor readout in cell-based measurements, three-dimensional diffusion simulations were performed on an adherent cell on a sensor-covered surface (Figure 3). The cell functioned as a physical barrier that blocks diffusion. This geometry mimics experimental configurations in which substrate-bound sensors are positioned beneath adherent cells.^34^

**Figure 3:**
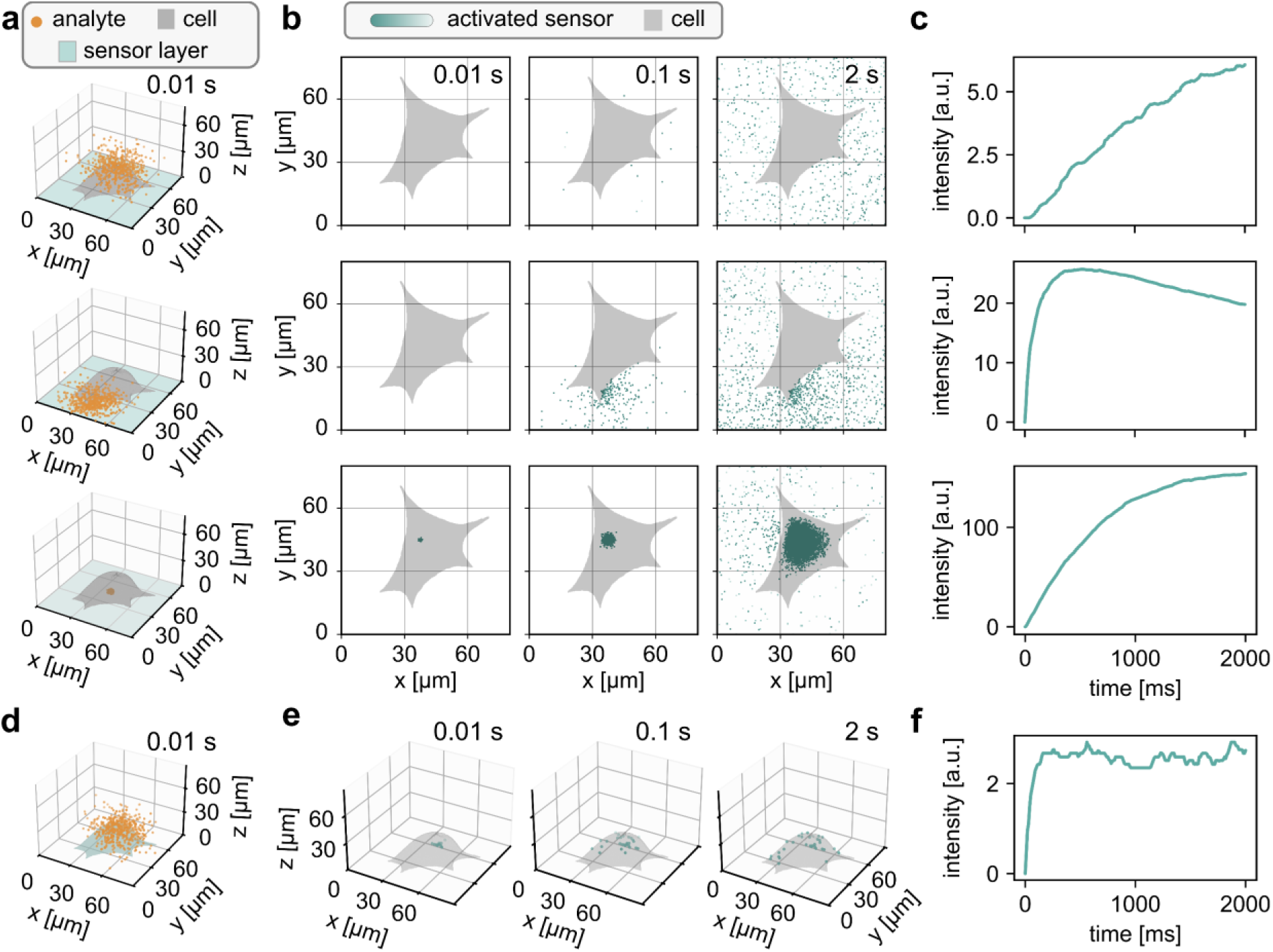
Simulation of analyte release by a cell from different locations. **a)** 3D diffusion simulation of analyte molecules at t = 0.01 s following release from different positions relative to a cell (gray). 10^7^ analyte molecules (orange) released from the top of the cell (top row), released from the side near the bottom (middle row) and released from underneath the cell (bottom row). The cell is positioned 100 nm above the bottom surface to mimic a typical adhesion scenario. The surface is covered by sensors (green). **b)** Top-down view: spatial distribution of sensor binding at t = 0.01, 0.1, and 2 s (left to right) for each release scenario. Each pixel (256×256 grid) represents one sensor with 30 binding sites. **c)** Time course of total bound molecules on the sensor surface for each release scenario. **d)** 3D diffusion simulation of analyte molecules at t =0.01 s following release from the top of the cell. The cell-membrane is covered by sensors (green). Each pixel has 1 binding site. **e)** 3D view of sensor-analyte complexes (=activated sensor) at t = 0.01, 0.1, 2 s. **f)** Time course of total bound analyte molecules over time in the whole area. Videos: S6-S8; Parameters: Table S3.

Three release scenarios were simulated – release from the top, from the side, and beneath the cell (Figure 3a). Sensor responses varied for these scenarios. Release from the top of the cell showed only minimal, delayed sensor binding (Figure 3c, top panel). Analyte molecules must diffuse around the cell to reach the sensors below. These results suggest that, in real cell experiments, sensor activation only minimally reports release from the top of the cell. Release from the side near the bottom triggered a strong initial response, followed by rapid decay (Figure 3c, middle panel). Analyte molecules quickly reached nearby sensors but then diffused in the extracellular space. Release from underneath the cell showed continuous signal increase during the observation period (Figure 3c, bottom panel). In this scenario, the diffusion rate 𝐷 underneath the cell was reduced to 10 % to account for hindered diffusion through cell-substrate junctions that anchor the cell to the surface. Note, that these sensor response pattern strongly depend on the sensor kinetics (SI, Figure S2).

To show a receptoŕs or a genetically encoded sensor’s perspective, membrane-bound sensors were simulated on the cell surface (Figure 3d-f). Analyte molecules were released from the top of the cell, which caused the most rapid response. Because they are positioned directly at the release site, they respond faster than substrate-bound sensors, whose signal depends on the diffusion distance between the release site and the sensor plane.

To evaluate sensor performance under realistic experimental conditions we aimed to interpret experimental data (Figure 4). In this measurement, the release of catecholamines from human neutrophilic granulocytes triggered by activated platelets was recorded using DNA functionalized (GT)_10_-SWCNT sensors following a previous publication.^51^ For the simulation, cells were placed above a sensor-covered substrate according to the outlines of the bright field images. The experimental data (Figure 4b) showed strong fluorescence changes under few cells as well as a general fluorescence increase everywhere.

**Figure 4:**
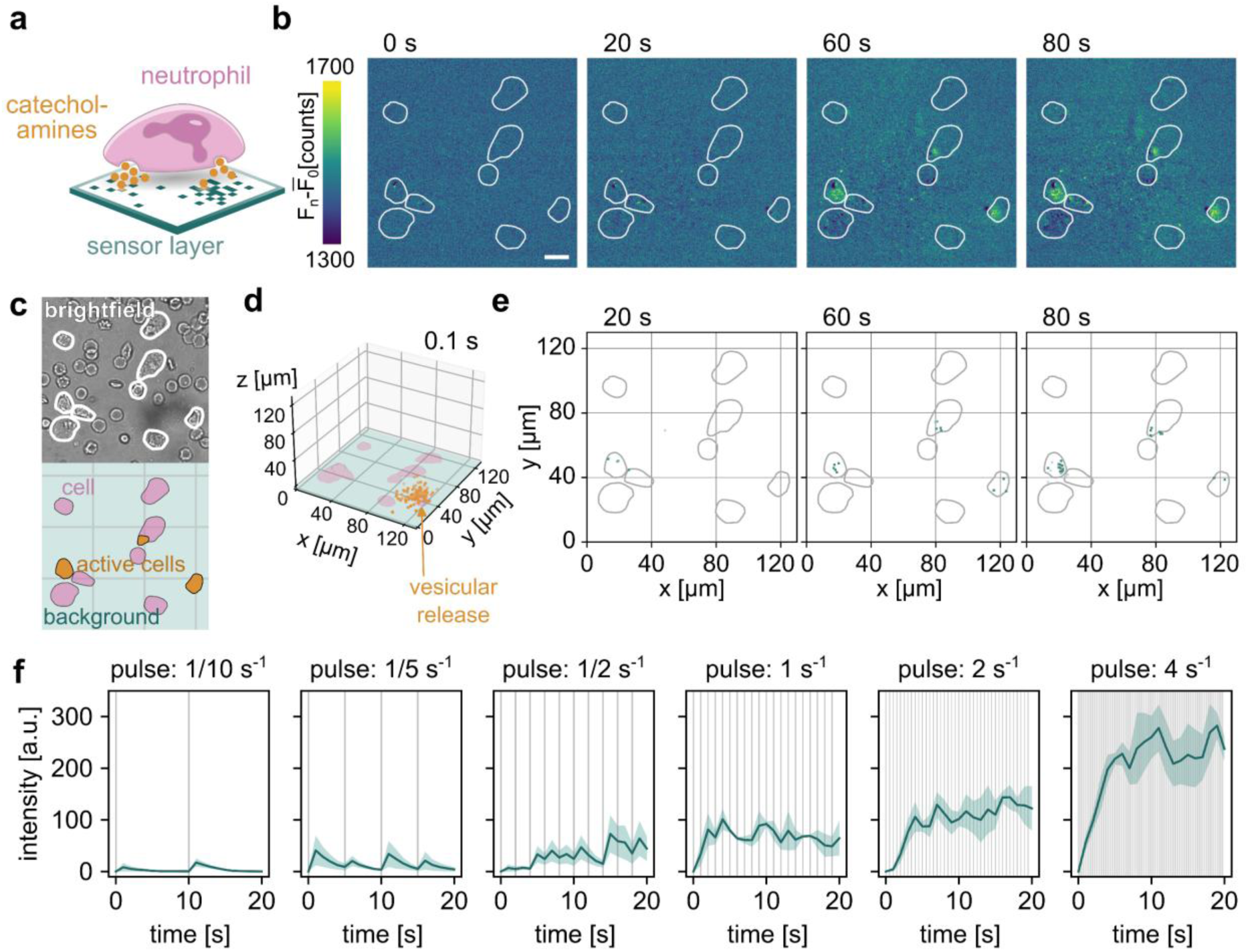
Simulation of paracrine signaling by immune cells. **a)** Human neutrophils release catecholamines after stimulation with activated platelets, which changes the fluorescence of nanosensors under the cells.^51^ **b)** NIR fluorescence images (fluorescence change) of DNA-SWCNTs sensors activated by catecholamines released from neutrophils at t = 0, 20, 60, and 80 s. White outlines indicate cell positions. Scale bar = 20 μm. **c)** Brightfield image (upper part) shows neutrophils surrounded by red blood cells. Input image for simulation (lower part). The sensors cover the entire bottom plane (green), cells are placed according to the brightfield image (pink), and release is restricted to three cells (orange). **d)** 3D spatial distribution of analyte molecules (orange dots) at t = 0.1 s and the first vesicular release. Six cells are modeled with a height of 20 μm and positioned 100 nm above the sensor surface. **e)** Simulation response of the scenario described in e, at t= 20 s, 60 s, 80 s. Exposure time = 1 s. Grey outlines indicate cell position. In total 3,300,000 molecules were released (3 cells x 130 vesicles x 33,000 molecules, released at different release/pulse rates), see also SI Figure S1**. f**) Substrate sensor response for different pulse/release rates. The shaded area shows the standard error (n= 5 simulations). Videos: VideoS9; Parameters: Table S4.

It probably reflects the different activation status of the cells or the release site and pinpoints to a huge heterogeneity. We thus simulated that three cells simultaneously released analyte molecules from their basal surface. Releases from upper cell surfaces were neglected because they contribute minimally to sensor detection. We simulated by 3 cells x 130 vesicles x 33,000 molecules (at timepoint 𝑡 = 0 𝑠) motivated by catecholamine numbers known from PC12 cells.^52^

The spatiotemporal distribution of sensor responses increased close to certain cells (Figure 4b). Additionally, the fluorescence signal increased gradually over time (Figure 4b,e). This is consistent with vesicles releasing catecholamines in successive bursts with decreasing frequency, rather than all molecules being released at once. In our simulation, 33,000 molecules were released per 130 vesicles and per active neutrophils, with release intervals speeding up from 8 s to 0.25 s (see SI Figure S1). The release interval profile was determined by reverse engineering the observed SWCNT fluorescence dynamics. Known biological parameters (diffusion, sensor kinetics, vesicle count, and molecule count per vesicle) were held fixed throughout, ensuring the simulation remained within physiologically validated bounds.

To test the limits of resolution, we simulated steady-state vesicular pulses across a range of frequencies from one release every 10 s up to four releases per second (Figure 4f). At low frequencies (≤ 1/5 s⁻¹), each release event appears as a discrete peak with full baseline recovery. Once the pulse rate exceeds 1/2 s⁻¹, however, peaks begin to merge and the signal no longer returns to baseline between events. At the highest frequencies, individual release events become indistinguishable in the time trace. Resolving it in space became also more challenging (Figure 4e). This frequency-dependent behavior has a practical consequence for data interpretation: a smooth, continuously rising signal does not necessarily mean catecholamines were released all at once. It may simply mean that vesicle fusion events occurred faster than the sensor could resolve them individually. The results also show that using our simulation it is possible to distinguish different biological release scenarios.

## Conclusion

The presented simulation framework addresses a major challenge in the quantitative interpretation of fluorescence microscopy data. Fluorescence images of biosensors do not directly report the local analyte concentration. They report the number of occupied binding sites of the sensor and how this translates into a fluorescence change. Additionally, the time course is shaped by sensor kinetics, geometry, and analyte concentration. This distinction is frequently overlooked in the interpretation of biosensor data. Only when association rate constants (𝑘_𝑜𝑛_) and dissociation rate constants (𝑘_𝑜𝑓𝑓_) are fast relative to the timescale of concentration changes or under equilibrium conditions fluorescence changes can directly be interpreted as concentration changes.^39^

Sensors with identical association constants 𝐾_𝐷_ but different rate constants produce qualitatively different images.^39^ Slow rate constant sensors integrate signals over time and cannot resolve fast phasic events. Moreover, slow 𝑘_𝑜𝑓𝑓_ rate constants reduce the pool of freely diffusing analyte molecules, as occupied sensors act as sinks and thus similar to transporters such as DAT. Therefore, sensors distort not only the local concentration but also diffusion. Together, these findings indicate that sensor design should prioritize the optimization of individual rate constants, not thermodynamic affinity alone. Embedding binding kinetics explicitly into the interpretation process has been shown to recover true analyte concentrations from fluorescence data with high fidelity.^53^ These factors also affect the fluorescence traces obtained from membrane-localized genetically encoded sensors compared to extracellular substrate-bound sensors (e.g. SWCNT-based). Our simulation reveals how the response curves depend on the position/placement of the sensors and which performance can be expected.

Measuring experimentally rate constants requires in the optimal case single-sensor, single-analyte experiments that are technically demanding.^37^ In this context, our simulation can be used to predict the optimal rate constants necessary to resolve a specific biological scenario. Additionally, it can be used to solve the inverse problem: Interpretation of a given experimental data set as outlined in Figure 4. In principle, different release scenarios can be simulated, and the simulated image series is compared to the experimental image series. Then, the difference is computed and minimized to predict the most likely scenario. This way one could narrow down the scenarios (release location, quantal size, spatiotemporal pattern). Overall, this simulation framework is broadly applicable to predict chemical signaling and its observables (fluorescence). Additionally, it provides quantitative insights to assess if a certain sensor technology can resolve biological questions and distinguish biological scenarios.

## Materials and Methods

### Simulation Algorithm Implementation

Molecular diffusion and binding simulations were implemented using Python with PyTorch, NumPy, and Matplotlib libraries. After acceptance the code will be made public on github.

### Diffusion Model

The diffusion process was simulated by a random walk in both 2D (n x n [pixel] or 3D (n x n x n [pixel]) environments. Each particle’s movement in one dimension is updated based on:

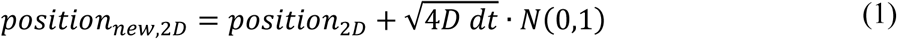

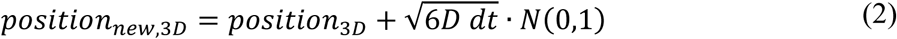

Here, d𝑡 represents the time interval and 𝐷 is the diffusion coefficient. 𝑁(0,1) represents a random number drawn from a standard normal distribution with mean 0 and standard deviation 1.

### Diffusion barriers

The framework uses reflecting boundary and thus all molecules stay in the n x n (x n) volume of the simulation. A binary PNG file can be provided to specify additional obstacles such as cell bodies. Each pixel is mapped onto the spatial grid and treated as either as free space or blocked region (impermeable barrier). Molecules what would enter barrier during diffusion are reflected by a negative step. If this reflection would also violate a boundary condition the molecule remains at its original position.

### Release Events

Particles can be initialized at different locations, in the middle of the area ([n/2, n/2] or [n/2, n/2, 0]) or in relation to a given cell pattern. Again, the area can be proved as a binary PNG file. Particles can be initialed on the edge of the cell, or in case of 3D also on top or bottom. In a uniform random distribution or in x different localized seeding, resembling vesicles. Also, the initialization can be a singular event in the beginning or distributed over time, in multiple release events.

### Removal Process - Transporter

Spatial sinks are defined by binary PNG masks and represent transporter-lined membrane surfaces. Uptake capacity is set by 𝑘_max_, the total number of transporters, with the counter initialized at 𝐶(0) = 𝑘_max_ all transporters free. At each time step, 𝐶(𝑡) evolves as in equation (3).

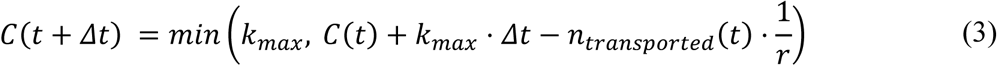

where 𝑟 is the transporter turnover rate 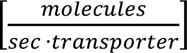 and *𝑛_𝑡𝑟𝑎𝑛𝑠𝑝𝑜𝑟𝑡ed_* (𝑡) is the number of molecules absorbed at step 𝑡.

### Surface Sensor and Binding Kinetics

The code includes a general reaction module for surface binding, used to compute sensor occupancy signals based on the local particle concentration. Sensors are distributed uniformly on the whole surface (2D) or on the bottom of the volume (3D). The sensors population can be limited to png-specific areas.

Each sensor binds with the on-rate constants 𝑘_𝑜𝑛_ and off-rate constants 𝑘_𝑜𝑓𝑓_.

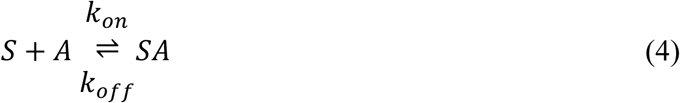

The binding probability 𝑝_𝑏𝑖𝑛𝑑_ and unbinding probability 𝑝_𝑢𝑛𝑏𝑖𝑛𝑑_ for every sensor 𝑖 can be described by equation (5) and (6).

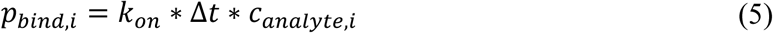

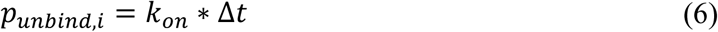

Here, Δ𝑡 is the simulated time interval and 𝑐_𝑎𝑛𝑎𝑙𝑦𝑡𝑒,𝑖_ is the analyte concentration at the position of sensor 𝑖. Sensors can have multiple binding side, here 30 binding sides per sensor. Binding events were filtered to follow a Poisson distribution, so there is only bound and unbound and no stated in between.

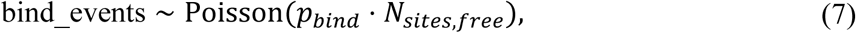

When bound the particles cannot diffuse. After unbinding, the particles can diffuse again.

### Isolation of human neutrophils from whole blood

Written informed consent was obtained from all blood donors prior to sample collection. Neutrophils were isolated from whole blood of healthy donors using the EasySep™ Direct Human Neutrophil Isolation Kit (Stemcell technologies, Vancouver; Canada). In this process, 7.5 ml of human blood is collected into a sterile K3 EDTA S-Monovette® (Sarstedt, Nümbrecht, Germany) and allowed to cool to room temperature (RT). Next, 3-4 ml of the blood is transferred to a 15 ml Falcon tube (Sarstedt, Nümbrecht, Germany). To this, 50 µl/ml of isolation cocktail and 50 µl/ml of rapid spheresTM are added, followed by a 5-minute incubation at RT. The sample is then brought to a total volume of 10 ml using PBS with 1mM EDTA, inverted for mixing, and placed in a magnetic field for 5 minutes. The supernatant is transferred to a new Falcon tube while in the magnet, with the same amount of rapid spheresTM added again, followed by another 5-minute incubation outside the magnet and then in the magnet. This process is repeated once more, with the supernatant being transferred to a new Falcon tube while in the magnet. The isolated cells are then centrifuged for 5 minutes at 100 g, the supernatant is discarded, and the cell pellet is carefully resuspended in 1 ml of Roswell Park Memorial Institute Medium 1640 (RPMI) for further use. The study was approved by the Ethics Committee Westfalen-Lippe (approval number 2021-657-f-S).

### Isolation of human platelets from whole blood

Platelets were isolated from citrated whole blood of healthy donors under the same ethical approval (2021-657-f-S) and with written informed consent. Whole blood was centrifuged at 259 × g for 30 min to obtain platelet-rich plasma (PRP). The PRP was further centrifuged at 259 × g for 10 min. Prostaglandin E1 (PGE1, final concentration 6.6 µM) was added to the PRP, which was then centrifuged at 10,000 × g for 30 s. The platelet pellet was gently resuspended in Tyrode’s buffer (pH 6.2) supplemented with PGE1 (5 µM) and centrifuged again at 10,000 × g for 30 s. After an additional wash under the same conditions, platelets were resuspended in Tyrode’s buffer (pH 7.4) and maintained at 37 °C until use.

### Neutrophil–platelet interaction assay

(GT)_10_-functionalized SWCNTs were immobilized on glass surfaces by incubation with a 4 nM suspension overnight at 4 °C. For control experiments, thrombin (1 U/mL; Sigma-Aldrich, USA) was applied either to blank SWCNTs or to SWCNTs bearing adhered platelets. For interaction assays, neutrophils (1 × 10⁶ cells/mL in HBSS) were seeded on SWCNT-coated surfaces and incubated for 5 min before addition of platelets, followed by another 5 min incubation. Measurements were initiated by adding thrombin (1 U/mL) after 10 s. Imaging was conducted using a 100× objective (UPLSAPO100XS, Olympus, Japan), with 561 nm laser excitation (120 mW) and 200 ms exposure at 5 frames per second. Drift was corrected in ImageJ using linear subtraction.

## Diversity, equity, ethics, and inclusion

This study was approved by the Ethics Committee Westfalen-Lippe (approval number 2021-657-f-S). Before donating blood, fully informed consent of each donor was obtained.

AI assistance: Perplexity, powered by Claude Sonnet 4.5, was used to improve clarity and readability of the manuscript text. All scientific content, data analysis, and interpretations remain the work of the author.

---

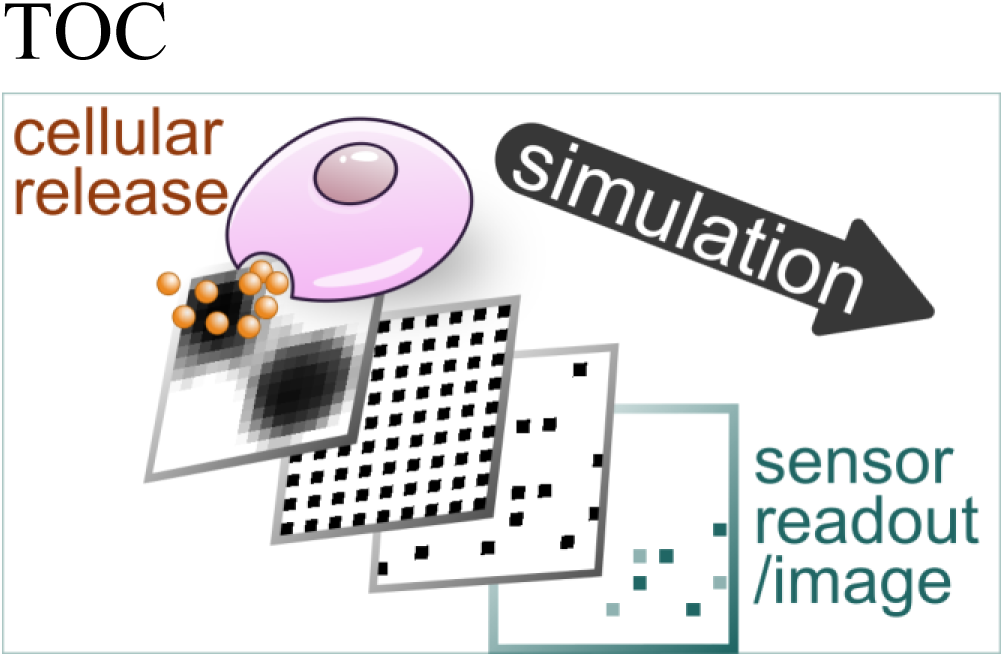

