## Supplementary material for "Simulation of neurotransmitter release and its imaging by fluorescent sensors": SI

### Supporting Information

Juliana Gretz[1], Jennifer M. Mohr[1], Bjoern F. Hill[1], Valeriia Andreeva[1], Luise Erpenbeck[2], Sebastian Kruss[1,3,*]

[1] Department of Chemistry and Biochemistry, Ruhr University Bochum; 44801 Bochum, Germany

[2] Department of Dermatology, University Hospital Münster; 48149 Münster, Germany

[3] Fraunhofer Institute for Microelectronic Circuits and Systems, 47057 Duisburg, Germany

Table S1: Parameters Figure 2 a-c

| Diffusion style | 2D | Source |
|---|---|---|
| Diffusion coefficient | $6{,}05 \cdot 10^{-6} cm^2 s^{-1}$ | (Trouillon et al., 2013) |
| k_on | $k_{on} = \frac{1}{\tau_{on} \cdot c} = \frac{1}{23ms \cdot 5 \cdot 10^{-6}M} = 8.7 \cdot 10^6 M^{-1} s^{-1}$ | (Kagiampaki et al., 2023) |
| k_off | $k_{off} = \frac{1}{\tau_{off}} = \frac{1}{194ms} = 5.15\ s^{-1}$ | (Kagiampaki et al., 2023) |
| Concentration | 3 vesicles with 30.000 molecules each | (Omiatek et al., 2013) |
| Size [real] | 4 µm x 4 µm | (Milo et al., 2019) |
| Size [simulation] | 256 pixel x 256 pixel | |
| Binding sides | 1 per pixel | |
| Simulation | Every 1 µs for 50 ms total | |

Related Video: VideoS1

Table S2: Parameters Figure 2 d-e

| Diffusion style | 2D | Source |
|---|---|---|
| Diffusion coefficient | $6{,}05 \cdot 10^{-6} cm^2 s^{-1}$ | (Trouillon et al., 2013) |
| k_on | $k_{on} = 1 \cdot 10^5 M^{-1} s^{-1}$ | (Gretz et al., 2025) |
| k_off | $k_{off} = 1\ s^{-1}$ | (Gretz et al., 2025) |
| Concentration | 3 vesicles with 3.000 molecules each | (Omiatek et al., 2013) |
| Size [real] | 4 µm x 4 µm | (Milo et al., 2019) |
| Size [simulation] | 256 pixel x 256 pixel | |
| Binding sides | 3 per pixel | |
| DAT | 100 dopamine transporters<br>Uptake: 2 molecules per seconds | (Prasad & Amara, 2001) |
| Other transporter | 100 dopamine transporters<br>Uptake: 200 molecules per second | |
| Simulation | Every 0.01 ms for 2 seconds total | |

Related Video: VideoS2-S4

Table S3: Parameters Figure 3

| Diffusion style | 3D | Source |
|---|---|---|
| Diffusion coefficient | $6{,}05 \cdot 10^{-6} cm^2 s^{-1}$<br>$6{,}05 \cdot 10^{-7} cm^2 s^{-1}$ underneath cell | (Trouillon et al., 2013) |
| k_on | $k_{on} = 1.3 \cdot 10^4 M^{-1} s^{-1}$ | (Gretz et al., 2025) |
| k_off | $k_{off} = 0.4\, s^{-1}$ | (Gretz et al., 2025) |
| Concentration | 1 vesicle with 10,000,000 molecules | |
| Size [real] | 80 µm x 80 µm | |
| Size [simulation] | 256 pixel x 256 pixel | |
| Binding sides in b<br>Binding sides in e | 30 per pixel<br>1 per pixel | (Nalige et al., 2024) |
| Simulation | Every 1 ms for 2 s total | |
| Volume for sensor recogintion | Area of sensor * 100 nm | |

Related Video: S5-S8 (note that diffusing molecules (orange dots) do not represent all simulated molecules. The number was reduced for visibility.)

Table SI4: Parameters Figure 4

| Diffusion style | 3D | Source |
|---|---|---|
| Diffusion coefficient | $6{,}05 \cdot 10^{-6} cm^2 s^{-1}$<br>$6{,}05 \cdot 10^{-7} cm^2 s^{-1}$ underneath cell | (Trouillon et al., 2013) |
| k_on | $k_{on} = 1.3 \cdot 10^4 M^{-1} s^{-1}$ | (Gretz et al., 2025) |
| k_off | $k_{off} = 0.4\, s^{-1}$ | (Gretz et al., 2025) |
| Concentration | 3 cells with 130 vesicles with 33,000 molecules each | (Omiatek et al., 2013) |
| Size [real] | 130 µm x 130 µm | according to experimental conditions |
| Size [simulation] | 256 pixel x 256 pixel | |
| Binding sides | 30 per pixel | (Nalige et al., 2024) |
| Simulation | Every 5 ms for 90 s total | |
| Volume for sensor recognition | Area of sensor * 100 nm | |

Related Video: S9 (note that diffusing molecules (orange dots) do not represent all simulated molecules. The number was reduced for visibility.)

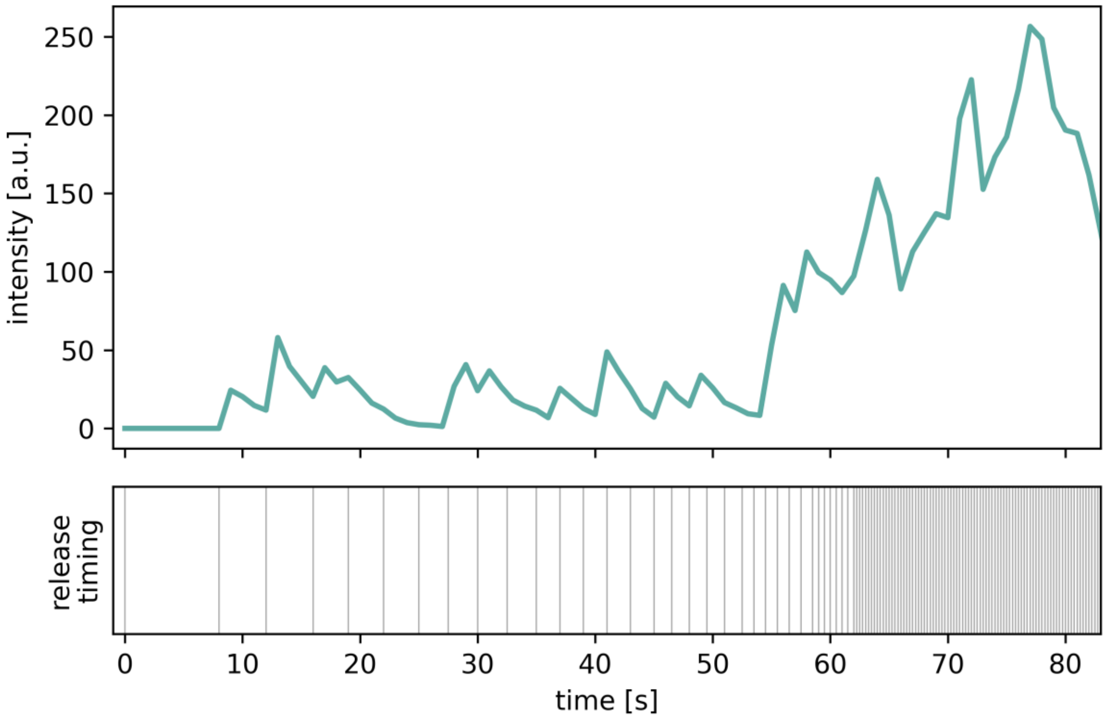


**Figure S1:** Vesicular release timing pattern in Figure 4e. The pulse frequency increases from 1/8 $s^{-1}$ to 4 $s^{-1}$.

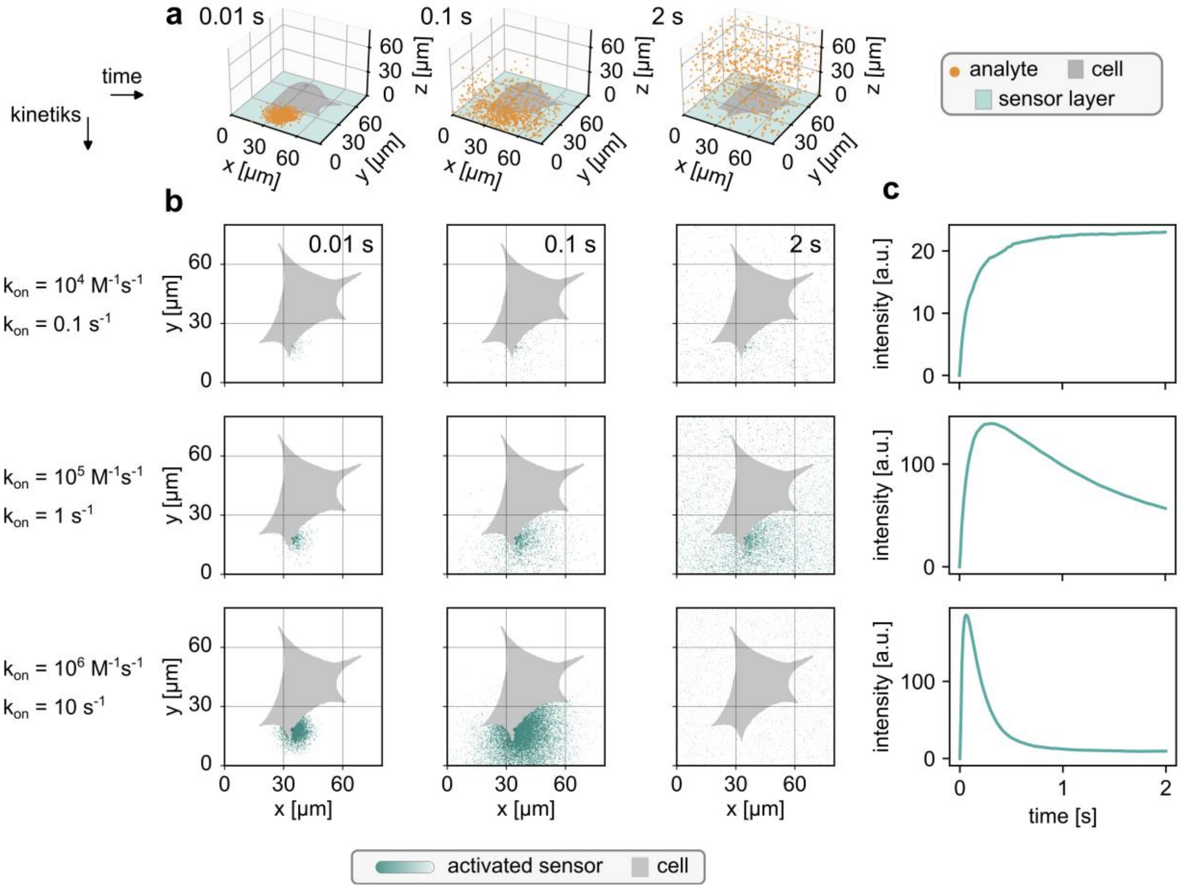


Figure S 2: **Simulation of analyte release with different sensor kinetics. a)** 3D diffusion simulation of analyte molecules at t = 0.01, 0.1 and 2 s following release from the side near the bottom. **b)** Top-down view: spatial distribution of sensor binding at t = 0.01, 0.1, and 2 s (left to right). Every row was simulated with different rate constants: top ($k_{on} = 10^4 M^{-1} s^{-1}$ and $k_{off} = 0.1\ s^{-1}$) , middle ($k_{on} = 10^5 M^{-1} s^{-1}$ and $k_{off} = 1\ s^{-1}$) , bottom ($k_{on} = 10^6 M^{-1} s^{-1}$ and $k_{off} = 10\ s^{-1}$) Each pixel (256×256 grid) represents one sensor with 30 binding sites. **c)** Time course of total bound molecules on the sensor surface for each release scenario.
